# Fast Batch Alignment of Single Cell Transcriptomes Unifies Multiple Mouse Cell Atlases into an Integrated Landscape

**DOI:** 10.1101/397042

**Authors:** Jong-Eun Park, Krzysztof Polański, Kerstin Meyer, Sarah A. Teichmann

## Abstract

Increasing numbers of large scale single cell RNA-Seq projects are leading to a data explosion, which can only be fully exploited through data integration. Therefore, efficient computational tools for combining diverse datasets are crucial for biology in the single cell genomics era. A number of methods have been developed to assist data integration by removing technical batch effects, but most are computationally intensive. To overcome the challenge of enormous datasets, we have developed BBKNN, an extremely fast graph-based data integration method. We illustrate the power of BBKNN for dimensionality-reduced visualisation and clustering in multiple biological scenarios, including a massive integrative study over several murine atlases. BBKNN successfully connects cell populations across experimentally heterogeneous mouse scRNA-Seq datasets, which reveals global markers of cell type and organ-specificity and provides the foundation for inferring the underlying transcription factor network. BBKNN is available at https://github.com/Teichlab/bbknn.

## 1 Introduction

The past few decades have seen a constant increase in the amount of data that can be harvested from single transcriptomics experiments^1,2^, culminating in the development of single cell RNA-Seq^3^. The single cell ‘resolution revolution’ in transcriptomics has enabled the analysis of different cell populations within a sample, instead of measuring an average across a heterogeneous input as for traditional bulk transcriptomics^4^. While various technological advances such as the introduction of Unique Molecular Identifiers (UMIs) have continued to improve gene expression quantification^5^, increase in throughput via innovations such as droplet microfluidics^6^ and combinatorial indexing^7^ have meant that the sizes of experiments and datasets have increased from e.g. 85 cells in 2011^8^, to atlases featuring over 100,000 cells^9^. These developments combined with continuously dropping sequencing costs and the fact that scRNA-Seq methods themselves are becoming cheaper to execute^10^, have led to the initiation of large-scale projects like the Human Cell Atlas^11^, emphasizing the urgent need for efficient computational approaches.

Given the variety of experimental methods generating vast amounts of data, efficient methods for aligning diverse data are invaluable for integrative analysis. A number of innovative batch correction approaches for scRNA-Seq have been proposed recently. This includes mnnCorrect^12^, which corrects expression against a reference batch based on the profiles of mutual nearest neighbours. Another example is Seurat’s CCA^13^, which merges cell types by performing CCA dimensionality reduction and aligning the resulting coordinate distributions across batches. Despite their merits, these methods often struggle with the exponential increase in sizes of data sets. The most efficient available approach is Scanorama^14^, which builds upon the ideas of mnnCorrect by identifying pairs of mutual nearest neighbours between the datasets, and using them to calculate a joint expression panorama. This method comes with its own form of computational limitations, requiring heavy RAM use for merging hundreds of thousands of cells.

Here, we present BBKNN (batch balanced k nearest neighbours) as a simple, fast and lightweight batch alignment method that integrates seamlessly into SCANPY^15^, a scalable Python scRNA-Seq analysis package. The output is immediately useable in downstream analyses, such as clustering^16^ and pseudotime inference^17^ along with UMAP^18^ and force-directed graphs^19^ for visualisation. Moving batch correction to the neighbour graph inference step allows for the creation of an extremely efficient algorithm with minimal computational requirements. Our comparative benchmarks show that BBKNN is one to two orders of magnitude faster than existing methods.

We demonstrate BBKNN’s utility in successfully merging data from two variants of the 10X Chromium^20^ droplet-based scRNA-Seq protocol and in integrating data from four pancreatic datasets stemming from different experimental setups^21–25^. Finally, we show how BBKNN aligns hundreds of thousands of cells from eight independent large-scale mouse scRNA-Seq data sets^4,9,26–31^. From this alignment, we identify intuitive developmental trajectories for mouse lineages, which we use as the framework to infer a transcription factor regulatory network for almost two thousand mouse transcription factors.

## 2 Results

### 2.1 Batch Balancing the Neighbourhood Graph

A common step in scRNA-Seq analysis is the identification of a neighbourhood graph, which is subsequently used for a number of downstream computational analyses, including clustering^16^ and pseudotime inference^17^, and visualisation methods such as UMAP^18^ and force-directed graphs^19^. The current default method for creating this graph within SCANPY is based on the approach proposed in UMAP. The graph concept is illustrated in Figure 1a: for each cell, its k nearest neighbours in principal component space are identified, and their Euclidean distances are then converted into connectivities with an exponential relationship. At this step, extra emphasis is placed on mutual neighbour cell pairs, with their two respective connectivities being replaced by their sum diminished by their product. However, when there is a batch effect present in the data, strong experimental variation can lead to cells being unable to form cross-batch biological connections.

**Figure 1:**
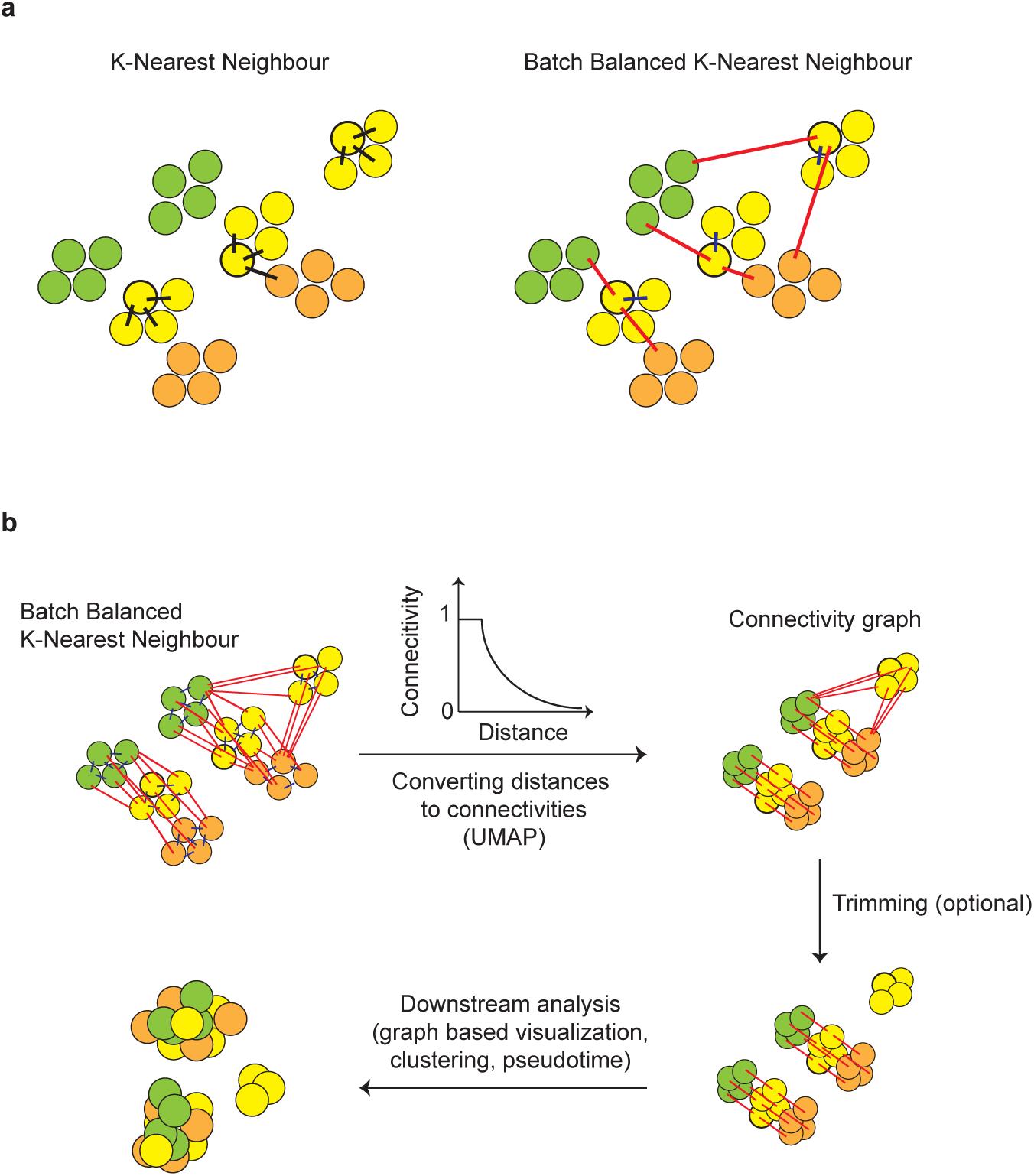
Workflow for single-cell data integration by BBKNN. **(a)** Identifying a cell’s neighbours by standard KNN as compared to the batch balanced counterpart in BBKNN. **(b)** The neighbour distance collections are then converted to exponentially related connectivities. BBKNN has an optional graph trimming step to weed out any erroneous connections between independent cell populations. The resulting connectivity graph can be used in downstream analyses such as clustering or UMAP visualisation.

Given the fact that neighbourhood graph construction involves identifying the nearest neighbours of each cell, and scRNA-Seq batch correction methods often make use of cross-batch neighbour information^12,14^, the procedure lends itself well to a very simple alteration that enables batch alignment. The BBKNN graph is constructed by defining the k nearest neighbours for each cell within each of the user-defined batches. This abstracted neighbour collection is then processed into graph connectivities in the same manner as when constructing a KNN graph, aligning the batches. In order to avoid aligning unrelated cells across batches when no equivalent cells exist in another batch, we limit the total number of edges for each cell. This prioritises mutual neighbour pairs and connections to similar cells across batches (Figure 1b). The resulting graph structure is immediately useable in the same broad range of downstream analysis options as the KNN graph. BBKNN also offers the option to compute approximate nearest neighbours, with run time scaling linearly and offering superior performance for datasets with hundreds of thousands of cells.

We illustrate this concept on simulated data as described in Supplementary Methods. The simulation confirms that BBKNN connects cells from a known shared population across experimentally different batches. It also demonstrates the use of the BBKNN trimming parameter to ensure that unrelated cells remain distinct, as shown in Supplementary Figure 1. Below, we focus on applying the algorithm to various biological datasets.

### 2.2 Merging PBMC Data from Droplet Protocol Variants

We first evaluate BBKNN in a biological context on two publicly available peripheral blood mononuclear cell (PBMC) samples, obtained using the 5*′*and 3*′*droplet protocols of 10X Chromium^20^. The 5*′*version was developed in response to community demand for accurate T and B cell receptor capture, allowing explicit VDJ reconstruction. Given that these methods capture different region of mRNA, we expect to see differences in gene expression quantification between cells profiled with 5*′*or 3*′*protocols. This is exactly what we observe: there is complete separation by experimental method when inferring a standard KNN neighbourhood graph (Figure 2a). However, upon BBKNN merging, this becomes supplanted by the various cell types integrating into unified clusters representing data from both experimental protocols (Figure 2b and 2c).

**Figure 2:**
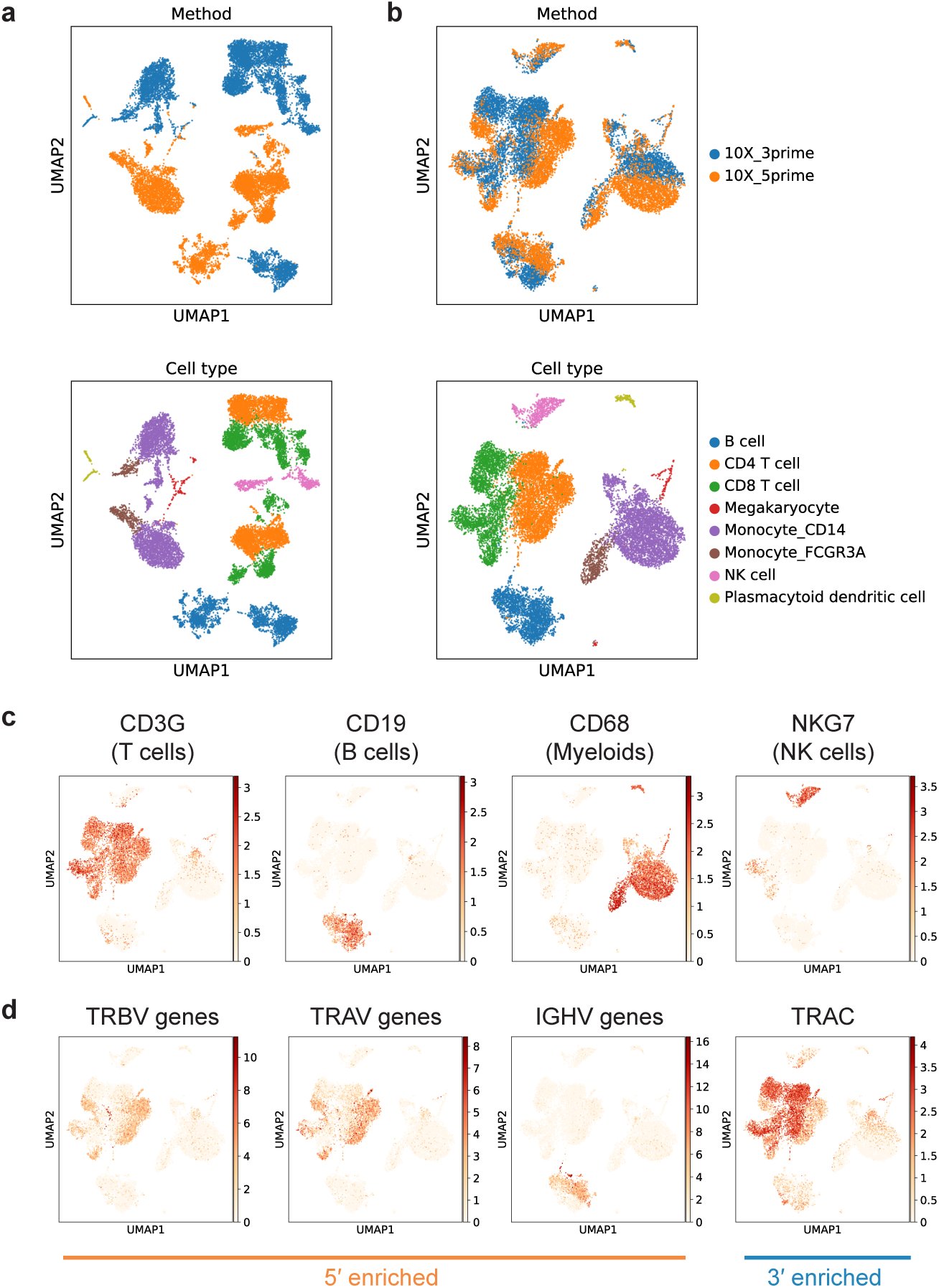
Integrating PBMC populations obtained via the 5*′* and 3*′* versions of 10X Chromium’s droplet technology. **(a)** Without any batch correction, the cells clearly split based on the experimental protocol. **(b)** Applying BBKNN successfully merges the cell populations, and recaptures cohesive subpopulation structures. **(c)** A selection of canonical markers identify the cell populations. **(d)** Residual heterogeneity within T and B cell populations stems from the differential capture efficiency of V and C gene segments of TCR and BCR genes between protocols.

Notably, a closer examination of the distribution of cells profiled by each protocol within the T and B cell clusters reveals genes expected to be technology-specific. For instance, the TRBV, TRAV and IGHV genes are captured by the 5*′*kit, while TRAC instead mainly appears in the cells from the 3*′*kit (Figure 2d). This is concordant with the fact that the detection efficiency of different regions of T and B cell receptors is the main difference between the two methods. As such, BBKNN performs excellent batch alignment for this data. BBKNN succeeds in allowing inference of a cluster structure that correctly captures the primary populations present in the data, while simultaneously retaining the protocol-driven biological differences in the relevant cell types.

### 2.3 Aligning Four Pancreatic Datasets from Diverse Technologies

Having demonstrated BBKNN’s ability to successfully merge cell populations from different droplet protocols, the algorithm was applied to a more diverse collection of publicly available pancreatic single cell data^21–24^. The experiments came from four independent studies, and were performed using a combination of both droplet^32^ and plate^33,34^ based methods. With around 15,000 cells in total between the four datasets, and a known shared biology captured in a set of standardised annotations^25^, the data provides a perfect testing ground to evaluate BBKNN. We compare its performance to the established batch correction methods mnnCorrect^12^, CCA^13^ and Scanorama^14^.

Figure 3a demonstrates a UMAP visualisation of the data without any attempt to remove the batch effect. A very clear separation can be seen based on the experiment of origin: while cells cluster together based on cell type within batches, corresponding cells across batches are widely scattered. The major cell types in the pancreatic islets are four endocrine cell populations: alpha, beta, gamma and delta. Additional smaller populations include stellate cells, mast cells and macrophages. An interesting point is that the gamma and delta pancreatic cells are already merged within the datasets at this stage.

**Figure 3:**
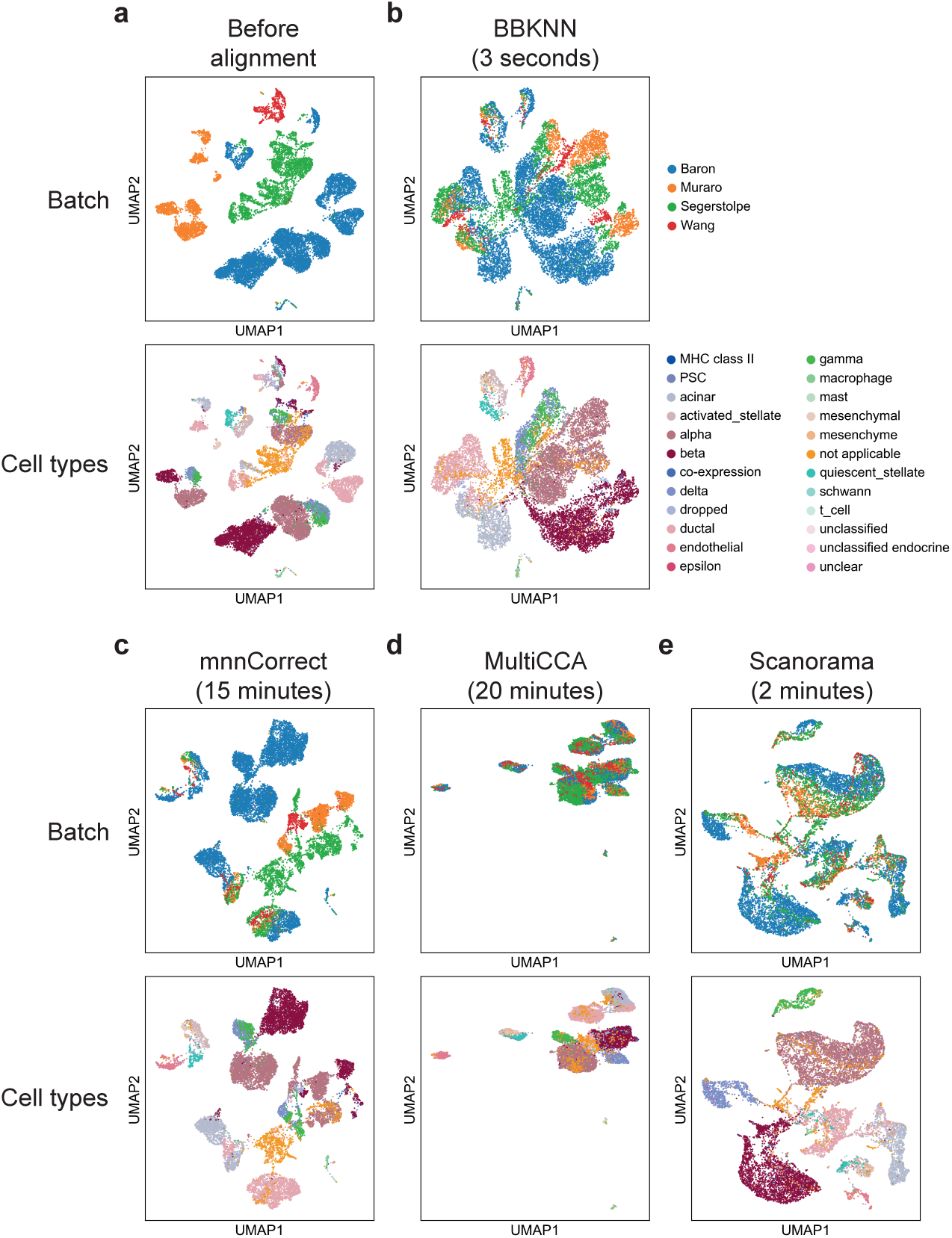
Applying BBKNN along with established batch correction methods to four experimentally diverse pancreatic datasets. **(a)** The data features large technical differences. The standardised cell annotations reveal the shared underlying biology. **(b)** BBKNN connects the cell populations in accordance with the annotation. **(c)** mnnCorrect retains residual technical differences in the data. **(d)** CCA mixes the batches well and correctly clusters the underlying cell types. **(e)** Scanorama performs to a similar standard, but is mildly challenged by some of the rare populations. When benchmarking the methods on a personal computer, BBKNN completes within three seconds, while the two methods implemented in R take upwards of 15 minutes and Scanorama takes 2 minutes.

After applying BBKNN (Figure 3b), all cell types are merged into the correct clusters, reflecting the information provided in the annotations. Examining the distribution of the experiment of origin reveals the batches to be far better mixed than in Figure 3a, although the stark experimental differences lead to a less homogeneous mixing than in the case of the PBMC droplets. The gamma and delta populations remain mixed, reflecting the fact that they were already intertwined to a good degree in the visualisation with no batch correction.

In comparison, the mnnCorrect method does not manage to merge cell clusters across experiments as successfully (Figure 3c), and a greater technological effect is conserved in the output. CCA merges the batches exceedingly well in Figure 3d, also reconstructing the underlying cell type structure to a similar standard as BBKNN while managing to separate the gamma and delta populations. Scanorama manages to mix the batches nearly as well as CCA (Figure 3e), and successfully reconnects the primary cell populations while splitting the gamma and delta cells. However, the quiescent stellate population, cohesively recaptured by all other methods, becomes fragmented. Additionally, a macrophage and mast cell cluster that is distinct in all other visualisations becomes absorbed by the main UMAP manifold. For this particular application, CCA produces the best aligned output, with BBKNN and Scanorama not far behind, offering trade-offs between batch integration and maintaining the distinct cell clusters. The level of batch integration by each of the methods is assessed via Shannon entropy (Supplementary Figure 2), with BBKNN outperforming mnnCorrect and obtaining results that are slightly less mixed than Scanorama and CCA.

In terms of compute time, there are large differences between the methods, with BBKNN by far the fastest. The benchmarking Jupyter Notebooks capturing this performance on a personal MacBook Pro with 16GB RAM and a four-core i7 processor are available as part of the GitHub repository. CCA took over 20 minutes to run for this data. The original R version of mnnCorrect took approximately 15 minutes, while a third party Python reimplementation^35^ was closer to 20. Scanorama took two minutes, while BBKNN took three seconds. Thus, BBKNN is at least 300 times faster in processing this data collection than the established R methods and 40 times faster than Scanorama. BBKNN remains one to two orders of magnitude faster than the other methods when applied to multiple simulated datasets, with an additional benchmarking of BBKNN’s neighbour detection algorithms against Scanorama for larger collections. BBKNN with approximate neighbour detection manages to perform an order of magnitude faster than Scanorama. Both evaluations are described in more detail in Supplementary Methods and shown on Supplementary Figure 3. These differences will become more marked as datasets continue to increase in size and heterogeneity, and users want to interact with datasets in a flexible and rapid manner. Therefore, we anticipate BBKNN’s fast and lightweight graph alignment method to become a popular tool for both individual users and in the context of databases and web servers.

### 2.4 Capturing a Developmental Trajectory Across Large-Scale Mouse Single-Cell Atlases

With BBKNN’s utility illustrated on two different well-studied biological scenarios, and its output falling in line with that of established batch correction methods, we set our sights on a larger dataset that would be computationally taxing for existing methods. Recent times have seen a veritable flood of murine scRNA-Seq atlases. We have collated eight of these, encompassing early embryo development, fetal, neonatal, and adult stages of mouse development, covering cells from at least 26 different mouse organs^4,9,26–31^. Some of the data sets are large-scale, organism-wide collections (Mouse Cell Atlas, Tabula Muris), while others focus on individual organs or cell types (embryo, thymus, kidney, brain and hematopoietic stem cells). Integrating the data leads to a collection of 267,690 cells, with a discernible split based on the dataset of origin.

Applying BBKNN to the data led to the datasets being well connected with each other, creating density radially spreading from the center in UMAP space. Both the uncorrected and processed versions of the complete atlas are presented in Supplementary Figure 4. However, due to the independent studies being of different scopes, some cell types (notably hematopoietic stem cells) were dominant in the structure due to sheer number differences. To overcome this, the dataset was down-sampled to guarantee more even cell population sizes, reducing the cell total to 114,600. This reduced-scale collection showcases a better balance between the constituent populations (Figure 4).

**Figure 4:**
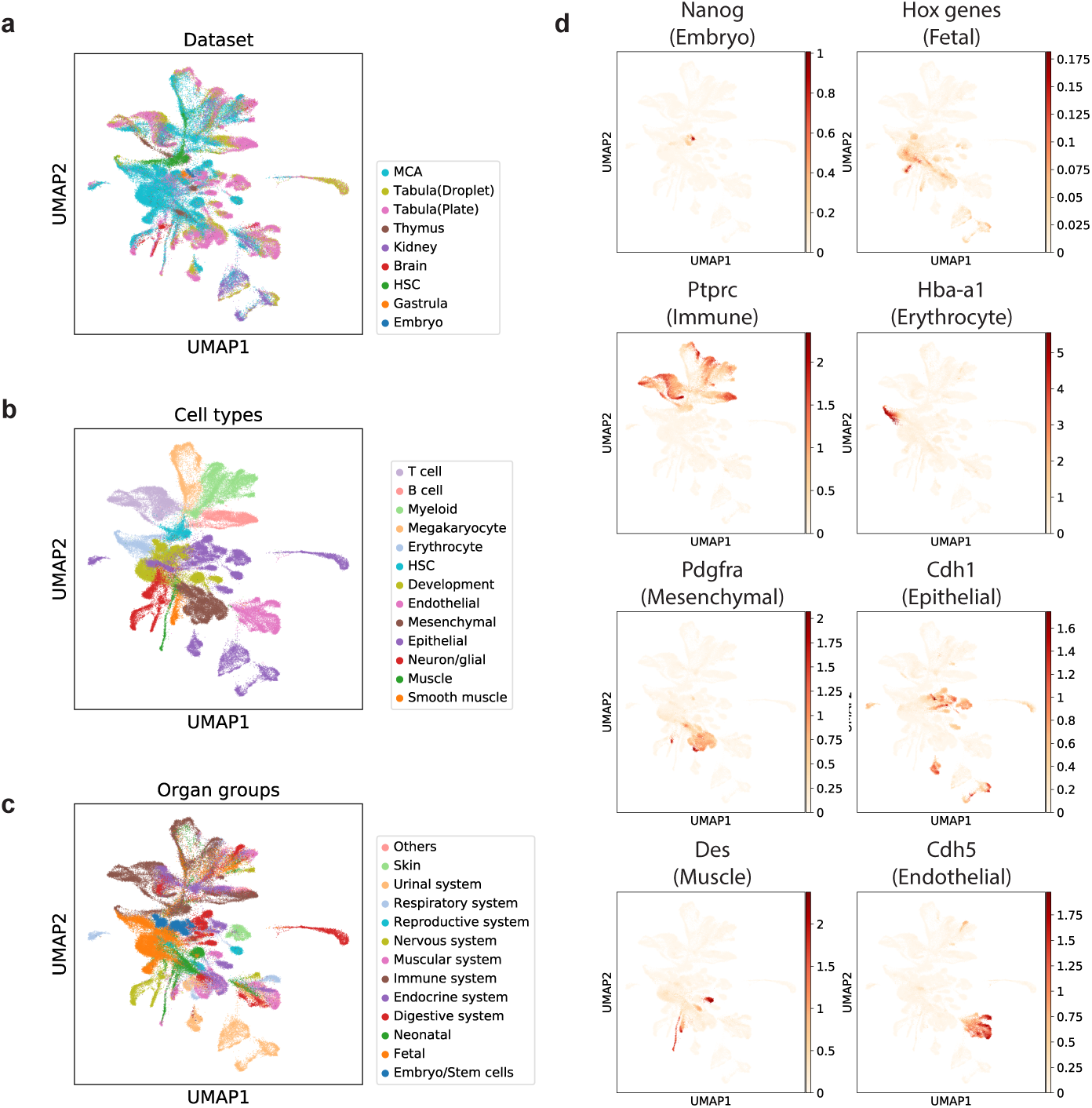
BBKNN-mediated integration of eight murine atlases with roughly 300,000 cells. **(a)** The algorithm merges data from eight independent atlases into a single cohesive entity. The cell populations have been down-sampled for visualisation purposes. The complete data can be seen in Supplementary Figure 4. **(b, c)** Embryonic cells are placed in the center of the cell space, and branch off to hematopoietic stem cells at the top and Hox-expressing fetal developing cells at the bottom. These progenitors give rise to mature immune and non-immune populations respectively. **(d)** A selection of canonical markers mirror the process and match the proposed annotation.

Cell populations were annotated by performing graph-based clustering^16^ and assessing canonical marker gene expression (Figure 4b and 4d). When examined along with the distribution of the atlas source (Figure 4a) and a simplified hierarchy of the reported organ of origin (Figure 4c), BBKNN appears to recapture a very intuitive development trajectory starting near the center of the UMAP manifold and working outwards. The root of the process are *Nanog*-expressing cells from developing embryos and embryonic stem cells, which lead to Hox-gene expressing fetal cells and hematopoietic stem cells. The hematopoietic lineage then branches into developed T cell, B cell, myeloid, megakaryocyte and erythrocyte populations. This developmental trajectory gains further support through the presence of early differentiating T and B cells, characterised by *Rag1* and *Igll1* expression at the junction with the stem cells (Supplementary Figure 5), underscoring the biological relevance of the relative positioning of the populations in the combined structure.

**Figure 5:**
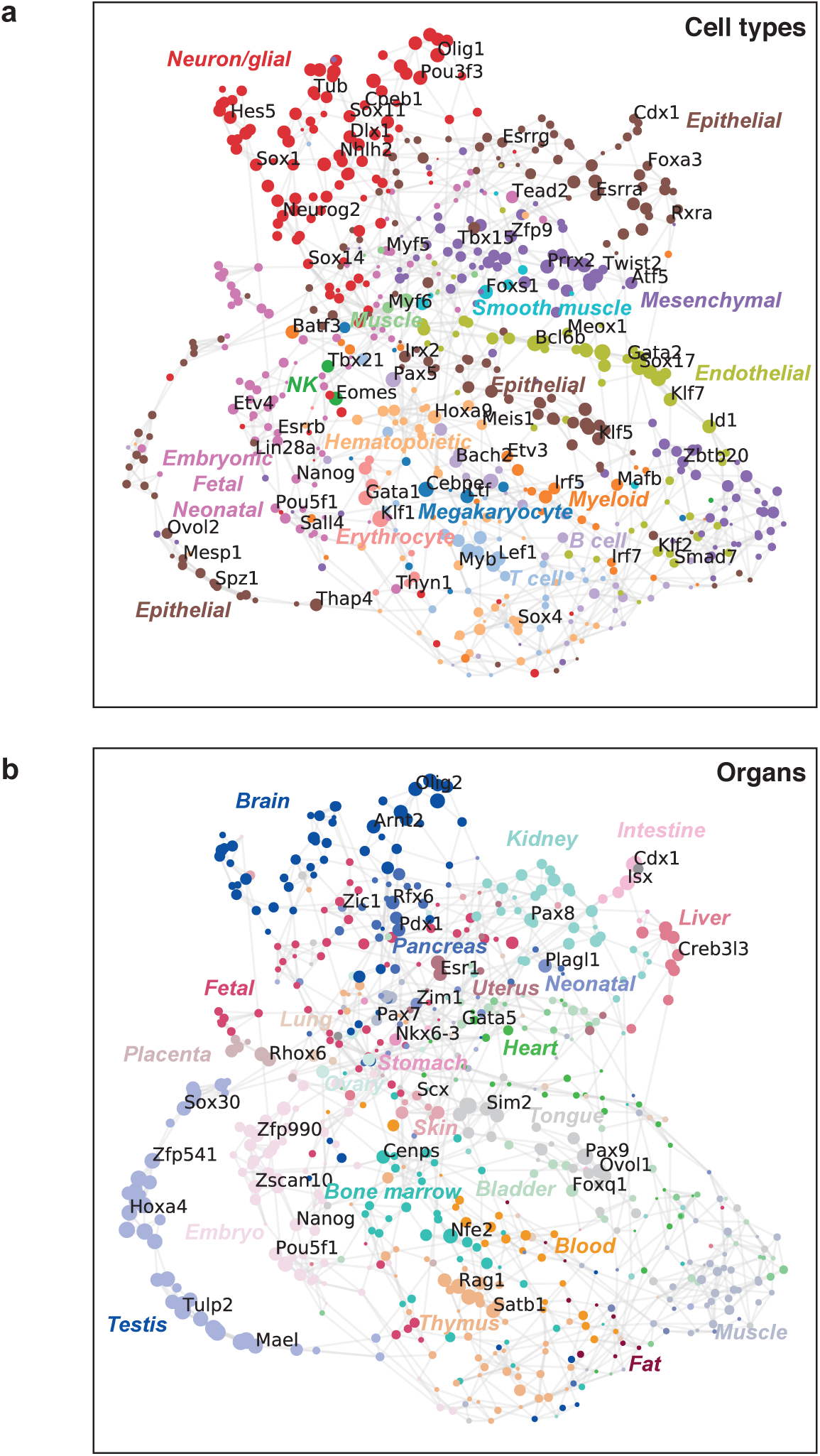
A transcription factor regulatory network inferred from the integrated murine atlases. Genes are coloured based on the most common **(a)** cell types and **(b)** organs expressing them, with the node size reflecting the specificity.

The opposite path captures the development of epithelial, mesenchymal, endothelial, muscular and neuronal cells. Most of these populations form tight, distinct clusters, with good blending of cells stemming from different organs. A notable exception was the distribution of epithelial cells, which revealed themselves to be highly divergent and organ specific. Numerous clusters of epithelial cells from the liver, lung, kidney and bladder are detached from the main body and are seen at different locations on the outskirts of the manifold, reflecting the specialised functionality of these tissue-specific subpopulations. A three-dimensional visualisation of the UMAP manifold can be seen as part of a dedicated exploratory Jupyter Notebook in the BBKNN GitHub repository, showcasing an even clearer radial trajectory from the centrally located stem cells.

### 2.5 Constructing a Comprehensive Mouse Transcription Factor Network

Given the biological cohesiveness of the BBKNN-corrected UMAP manifold of the murine atlases, which captured an intuitive developmental trajectory, we use it as a starting point to decipher the underlying transcriptional regulation. Network inference is a challenging task for single cell data due to sparsity. A proposed solution to this problem is expression imputation^36,37^, which we performed by creating ‘pseudo-cells’ as averages of the expression profiles of nearest neighbour cells in UMAP space. This pseudo-cell set was subsequently thinned according to the original UMAP coordinates of the averaged cells, resulting in a collection retaining the dynamic range of biological variation while offering denser expression matrices and reducing the total number of pseudo-cells by over tenfold.

We next selected a curated set of about 1700 murine transcription factors^38^ and analysed the correlation of gene expression patterns across the pseudocells. This provides a transcription factor UMAP manifold capturing a highly structured network with multiple modules (Figure 5). As this transcription factor network stems from single-cell resolution mouse gene expression atlases fully covering developmental and spatial diversity, this is one of the most comprehensive regulatory networks constructed to date.

A more detailed investigation into the exact functionality captured by the network was carried out by annotating each transcription factor node with information based on cell type and organ specificity. This was done by correlating each gene’s expression with the underlying distribution of the cell type and organ of origin for the constituents of each pseudo-cell. When examining the cell type annotation spread across the network (Figure 5a), there are clear modules highlighting transcription factors for each lineage. All members of the pluripotency core network (*Nanog, Pou5f1, Sall4, Esrrb, Lin28a*)^39^ form a cohesive cluster, and the cell type annotation accurately describes them as embryo-specific. This embryonic/fetal/neonatal branch of the network is connected to the center of the manifold, which serves as the branching point for modules specific to differentiated cell types. Known lineage marker genes are well captured in this representation (e.g. *Gata2* and *Sox17* for endothelial cells, *Hoxa9* and *Meis1* for hematopoietic stem cells, *Pax5* and *Bach2* for B cells, *Tcf7* and *Lef1* for T cells, *Twist2* and *Prrx2* for mesenchymal cells).

The heterogeneous, organ-specific behaviour of epithelial cells in the original manifold is mirrored in the regulatory network, with at least three distinct modules, each marked by specific transcription factors. The modules reveal themselves as very localised and specialised when examining the organ annotation (Figure 5b). The transcription factors with high organ specificity in the network are known master regulators of organ development (e.g. *Cdx1* and *Isx* for intestine, *Pdx1* for pancreas, *Rhox6* for placenta, *Nkx6-3* for stomach, *Pax8* for kidney and *Pax7* for muscle). Interestingly, transcription factors specific to neonatal tissue were enriched in imprinted genes (*Plagl1, Zim1, Zic1*)^40^, which is consistent with the notion that imprinted genes are crucial for regulating growth and development^41^. In contrast, the mesenchymal/endothelial module does not exhibit much organ specificity, reflecting the transcription factors’ general role in regulating these cell types across many organs. It would be interesting to investigate tissue specific nuances of these genes captured by the network.

The identification of established key regulators suggests that this analytical framework could serve as a useful data-mining tool for future integrative studies. We have created a dedicated exploratory Jupyter Notebook, featuring data from both the integrated cell space of Figure 4 and transcription factor network of Figure 5, and included it as part of the BBKNN GitHub repository.

## 3 Discussion

BBKNN is a very fast and lightweight batch alignment tool that can be used immediately with SCANPY^15^, a scalable Python single cell RNA-Seq package capable of handling over a million cells. The underlying idea is a simple alteration to the process of neighbourhood graph creation, replacing the standard KNN procedure with a batch balanced variant that identifies the closest neighbour for each cell within each of the experimental groups provided on input. This novel batch alignment concept is very quick to execute, making use of an efficient C implementation for neighbour identification^42,43^ and an approximate nearest neighbour alternative that scales better into huge datasets^44^. The resulting graph can be directly used as the input for downstream analyses such as clustering^16^ and diffusion pseudotime^17^, with compatible dimensionality reduction visualisation approaches including UMAP^18^ and force-directed graphs^19^.

We demonstrate BBKNN’s utility by applying it to a number of biological scenarios, and in each it was able to reconstruct the underlying shared cell populations. Examples are the four very disparate experimental setups of pancreatic data, and two biologically distinct protocols for PBMCs. Finally, we use BBKNN to propose an intuitive development trajectory in a landscape of hundreds of thousands of cells from murine atlases. The method was benchmarked against the established batch correction methods mnnCorrect^12^, CCA^13^ and Scanorama^14^ on the pancreatic data, yielding results of comparable quality, with run times one to two orders of magnitude faster on a personal computer.

A neighbourhood graph has a variety of downstream applications, and the choice of this format for batch alignment allows for easy deployment of a very fast and successful algorithm. At present, not all tools are equipped to work with neighbourhood graphs as input, with a notable example being SCANPY’s implementation of t-SNE^45^. However, our algorithm is perfectly compatible with UMAP, which is quickly gaining traction. Seurat^13^ has UMAP support, and the current development version of Monocle^46^ features trajectory inference within a UMAP-reduced space.

We demonstrate that BBKNN is able to integrate large and disparate data sets into a single structure by applying it to multiple large mouse single cell atlases, providing the biologic community with a valuable resource to gain insights into diverse fields ranging from developmental biology to tissue adaptation. The utility of the results is asserted by the murine cell atlas integration serving as a baseline for the inference of the underlying regulatory network, which captures the branching of embryonic transcription factors into various cell type and organ specific modules.

## 4 Methods

### 4.1 BBKNN

BBKNN was designed to slot into the spot occupied by neighbourhood graph inference in the typical SCANPY workflow, making use of PCA coordinates stored within the AnnData object. We replaced the standard KNN procedure with a variant that identifies nearest neighbours in each of the provided batches, creating a batch balanced version of the neighbour candidates which are subsequently processed into connectivities via the same function SCANPY’s neighbourhood graph creation uses. The neighbour identification itself is performed via `cKDTree`^42^ from `scipy.spatial`^43^ for the default Euclidean metric, with additional metrics supported by `KDTree` from `sklearn.neighbors`^47^ at reduced performance. BBKNN also supports annoy^44^, with its approximate nearest neighbour identification scaling better into large datasets. After the neighbour candidates are transformed into connectivities, the graph can be optionally trimmed to only feature a user-provided number of best connections for each cell to help ensure the independence of unrelated cell populations.

### 4.2 Seurat-Inspired SCANPY Workflow

The three biological scenarios were evaluated using a common analysis core, which shall be henceforth referred to as the Seurat-inspired SCANPY workflow. The steps of the analysis are normalising the data to 10,000 counts per cell, identifying highly variable genes, limiting the datasets to those genes only, log transforming the data, scaling it to unit variance and zero mean followed by PCA. At this stage, the established analysis identifies a regular KNN graph, but we also apply BBKNN in parallel. Both resulting AnnData objects are subsequently dimensionality-reduced with UMAP and are subjected to graph-based clustering.

### 4.3 Droplet PBMCs

The input data was downloaded from the 10X Genomics website. The exact 5*′*dataset was ‘PBMCs of a healthy donor 5*′*gene expression’, under Cell Ranger 2.1.0, under V(D)J + 5*′*Gene Expression. The exact 3*′*dataset was ‘8k PBMCs from a Healthy Donor’, under Cell Ranger 2.1.0, under Chromium Demonstration (v2 Chemistry). The core of the analysis was performed using the Seurat-inspired SCANPY workflow, with cell filtering on unique genes (between 500 and 7000) and total UMIs (above 2000) performed prior to the normalisation. Cells were annotated in the KNN object based on canonical markers, and this annotation was carried over and visualised in the BBKNN object.

### 4.4 Pancreatic Data

The data for the four different pancreatic experiments was downloaded in the form of homogeneously prepared `SingleCellExperiment` R objects featuring standardised annotations^25^. It was then processed with the Seurat-inspired SCANPY workflow. The standard neighbourhood graph analysis was compared to BBKNN, along with CCA and both the R and Python versions of mnnCorrect, with each being applied independently at their desired points in the analysis (as replacements to neighbourhood graph computation, PCA and prior to data scaling respectively). Scanorama was performed on raw data filtered to feature cells with a minimum of 600 unique genes, and its output was inserted at the PCA step of the Seurat-inspired SCANPY workflow. Run time benchmarking was performed on a personal MacBook Pro with 16GB RAM and a four-core i7 processor, and is captured in the Jupyter Notebooks hosted in the GitHub repository. Batch integration was assessed by performing Louvain clustering starting at zero resolution and increasing the parameter by a fixed step (0.005 for uncorrected, 0.01 for mnnCorrect/Scanorama, 0.05 for BBKNN and CCA), computing Shannon entropy as a measure of mixing and reporting it along with the cluster count. The clustering was terminated once over 15 clusters were identified for each correction method.

### 4.5 Murine Atlases

Both the droplet and plate data from Tabula Muris was downloaded from figshare, while all the other atlases were obtained from GEO. The dataset was then analysed with the Seurat-inspired SCANPY workflow (Supplementary Figure 4). To avoid cell type over-representation biases, the dataset was downsampled based on organ annotations provided within each atlas coupled with Louvain clustering of the complete object with a regular KNN graph – if a given atlas featured over 2,000 cells of a given organ, the cluster memberships of all of those cells were extracted. If any cluster featured over 500 cells from that organ in that atlas, those cells would be randomly sub-sampled to remove the excess. Additionally, the provided organ annotations were manually classified into a smaller set of more general labels for visualisation purposes, with both the original and simplified versions being contained in the provided AnnData objects. The Seurat-inspired SCANPY workflow was repeated after downsampling, with the addition of the creation of a three-dimensional UMAP manifold included in the exploratory Jupyter Notebook in the GitHub Repository.

For the transcription factor network, the cells were replaced with pseudocells created by averaging the expression of any selected cell’s 30 nearest neighbours in the three-dimensional UMAP. The space was divided into unit voxels. Each voxel featuring fewer than five pseudo-cells was emptied and remaining populated voxels were downsampled to 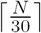 pseudo-cells, where *N* is the number of cells originally present in the voxel. The voxels had the proportion of each cell type and organ appearing among their cells calculated, assigning the values to their resulting pseudo-cells. These proportions were subsequently compared to gene expression with the Pearson correlation coefficient. The pseudo-cells were used to identify highly variable genes, the gene pool was filtered to feature high confidence transcription factors from AnimalTFDB3, the expression matrix was transposed and a UMAP manifold was proposed based on a cosine distance KNN graph of the genes. Edges were drawn between the nodes of the network where UMAP connectivity exceeded 0.25, node colouring was based on the most correlated cell type/organ annotation, and node size reflects the difference between its correlation coefficient and the next highest correlation coefficient in that category, accounting for specificity.

## Acknowledgments

The authors would like to thank Nadine Flinner for imputation assistance, Johan Henriksson for optimal trimming implementation inspiration, Oliver Stegle for suggesting the benchmarging computational experiment, Xi Chen, Mirjana Efremova, Kyowon Jeong, Tomás Gomes, Martin Hemberg and Roser Vento for feedback on the manuscript and all members of the Teichmann lab for helpful discussions.

## Author contributions

JP conceived the method, conducted the PBMC and murine analyses, created the figures and contributed to the manuscript. KP implemented the method and wrote the documentation, conducted the pancreas analysis/benchmarking and created the manuscript. KM and ST supervised the work and assisted in the writing of the manuscript.

